# Monkey Prefrontal Cortex Learns to Minimize Sequence Prediction Error

**DOI:** 10.1101/2024.02.28.582611

**Authors:** Huzi Cheng, Matthew V. Chafee, Rachael K. Blackman, Joshua W. Brown

## Abstract

In this study, we develop a novel recurrent neural network (RNN) model of pre-frontal cortex that predicts sensory inputs, actions, and outcomes at the next time step. Synaptic weights in the model are adjusted to minimize sequence prediction error, adapting a deep learning rule similar to those of large language models. The model, called Sequence Prediction Error Learning (SPEL), is a simple RNN that predicts world state at the next time step, but that differs from standard RNNs by using its own prediction errors from the previous state predictions as inputs to the hidden units of the network. We show that the time course of sequence prediction errors generated by the model closely matched the activity time courses of populations of neurons in macaque prefrontal cortex. Hidden units in the model responded to combinations of task variables and exhibited sensitivity to changing stimulus probability in ways that closely resembled monkey prefrontal neurons. Moreover, the model generated prolonged response times to infrequent, unexpected events as did monkeys. The results suggest that prefrontal cortex may generate internal models of the temporal structure of the world even during tasks that do not explicitly depend on temporal expectation, using a sequence prediction error minimization learning rule to do so. As such, the SPEL model provides a unified, general-purpose theoretical framework for modeling the lateral prefrontal cortex.

## 1 Introduction

Our response to environmental stimuli is often contingent upon our internal state or the behavioral goals we have established. Cognitive control provides this context dependence, allowing us to respond differently to environmental stimuli according to internal representations (Goldman-Rakic, 1987) of goals or rules, which depend on the lateral prefrontal cortex (Miller and Cohen, 2001). Cognitive control frees human behavior from rote or reflexive responses to stimuli, making our actions autonomous and self-regulated. Several neural correlates of cognitive control have been identified in primate lateral prefrontal cortex (PFC), including cells that encode objectively abstract parameters that could be used to govern how environmental inputs are converted into behavioral outputs, including cells that encode task rules (Wallis et al., 2001), cells that encode categories (Freedman et al., 2002; Crowe et al., 2013; Goodwin et al., 2012), cells that encode numerosity (Nieder and Miller, 2003; Ramirez-Cardenas et al., 2016), and cells that exhibit a property termed mixed selectivity by which different variables interact multiplicatively to influence firing rate (Rigotti et al., 2013).

The learning rule that prefrontal cortex uses to adapt neural information processing to changing cognitive demands is not known. This is an important question to answer because the computations that prefrontal cortex performs likely arise as a result of learning. Therefore, asking what computations prefrontal cortex performs and what learning rule prefrontal cortex uses to adapt computation to the environment are closely related questions. Greater understanding of the learning rule should provide deeper understanding of the functions of prefrontal cortex.

In this study, we develop a recurrent neural network model of prefrontal cortex that learns to perform a cognitive control task by minimizing sequence prediction error. At each time step in the task, the model utilizes patterns of activity in the hidden layer to predict the state of the environment (and the agent) in the next time step. Synaptic weights are adjusted on rewarded trials to improve the prediction by minimizing sequence prediction error. The task we trained the model to perform using this learning rule is a variant of the AX continuous performance task (AX-CPT) that has been widely used to measure cognitive control deficits that are associated with reduced activation of prefrontal cortex in schizophrenia (Barch and Ceaser, 2012; Barch et al., 2003; MacDonald, 2008; Lesh et al., 2013; MacDonald et al., 2005). We have recently used it to characterize neural activity in monkey prefrontal cortex (Blackman et al., 2023, 2016).

We reason that if a recurrent network is trained to perform a task that monkeys perform, and if neural dynamics in the recurrent network come to closely resemble neural dynamics in the monkey prefrontal cortex, then the learning rule used to train the recurrent network may closely approximate the learning rule implemented in the brain to train the prefrontal cortex. To evaluate the quality of the match between artificial and real neurons, we compare neural dynamics in our recurrent neural network simulation to neural dynamics directly observed in monkey prefrontal cortex while the real and artificial agents perform the same cognitive control task. This allows us to evaluate the number of neural signatures of computation in the brain that the model can successfully generate. One such neural signature described previously is provided by ’switch neurons’ in prefrontal cortex that were found to respond to infrequent stimuli countermanding a habitual response. These neurons switched their stimulus preference between different epochs of the trial depending on which stimuli were frequent (instructing habitual responses), and which stimuli were infrequent (instructing counter-habitual responses, therefore requiring cognitive control) (Blackman et al., 2016, 2023). The recurrent network architecture and learning rule we have developed replicates this form of neural activity.

Various theoretical and computational models have attempted to account for the function and role of the lateral prefrontal cortex as representing rules (Miller and Cohen, 2001), and how those may be learned hierarchically (Alexander and Brown, 2015, 2018). Still, many of these models rely on degrees of hand-tuning various aspects of the architecture and may not generalize to other tasks without additional hand-tuning of the network architecture or the learning rules, or both. Moreover while some cells represent a single task feature neatly (Wallis et al., 2001), lateral PFC representations are often complex, with various combinations of features represented in individual cells that show mixed selectivity (Rigotti et al., 2013). Candidate computational neural models of lateral PFC should be able to learn a variety of tasks autonomously and reproduce non-trivial aspects of learning, behavior, and neurophysiological cell types and representations.

As a starting point, the free energy principle (Friston, 2010) proposes conceptually that the brain minimizes prediction error, either by learning to better predict the world, or by acting to minimize the difference between intended and actual outcomes (Friston et al., 2011; Zarr and Brown, 2023). There is empirical support for the notion that monkey prefrontal cortex encodes sequences (Bellet et al., 2021; Averbeck et al., 2006), as well as errors in sequence prediction as it relates to reward (Oemisch et al., 2019). However, how such signals are used to govern learning in prefrontal cortex is not fully understood. Our model considers errors in sequence prediction that extend beyond rewards to entail more global representations of environmental state (Gläscher et al., 2010). This allows us to ask the question whether minimizing errors in state prediction may give rise to neural responses like those found in prefrontal cortex. We show below that a model trained to predict the state of the world in the next time step in a manner consistent with the free energy principle can account for a surprisingly broad set of findings in the lateral PFC. For this, we first require a computational principle and then a set of data to constrain the model development.

We approach this problem with a simple premise: **lateral prefrontal cortex is essentially a deep recurrent network that learns to predict the next state of the world moment by moment, with prediction errors as both inputs and training signals**.

The model simulates time at high resolution. Early cognitive models using deep learning have simulated the fundamental unit of time as a task trial (McCloskey and Cohen, 1989), which may be several seconds long. However no such artificial task delineation exists in the brain. Rather, neurons process stimuli and outputs millisecond by millisecond. This real-time approach to modeling provides a way to simulate a wide variety of cell type representations (Brown et al., 2004; Alexander and Brown, 2011).

With our approach, the model learns to store only what it needs to avoid prediction errors. The byproduct is a sufficient basis of representations for guiding temporally extended behaviors. The model will also clarify why PFC is mainly activated initially during behavior, and once a task is learned, PFC activity is not as strong (Chen and Wise, 1995).

Here, we develop a novel recurrent neural network model of prefrontal cortex, called the Sequence Prediction Error Learning (SPEL) model. The model may unify ongoing studies of prefrontal neurophysiology with recent advances in deep learning in recurrent networks. Specifically, we show that training a recurrent network using a sequence prediction error learning rule, similar to that employed by transformer architectures such as chatGPT (Vaswani et al., 2017), very readily replicates a broad array of neurophysiological findings obtained by recording neural activity from primate prefrontal cortex. Under this learning regimen, synaptic weights are adjusted to optimize the prediction about future sensory inputs and motor outputs that can be derived from the current activity state of the network at each time step throughout the trial. Implementation of this learning rule replicated much of the neural activity observed in monkey prefrontal cortex, including switch neuron activity, which was revealed to encode sequence prediction error in the model.

## 2 Results

Macaque monkeys were trained to perform a working memory task called the DPX (dot pattern expectancy) task (Blackman et al., 2016), shown in Figure 1. Briefly, the monkeys had to hold a cue in working memory, then respond to a subsequent probe based on both the cue and probe identities.

**Figure 1:**
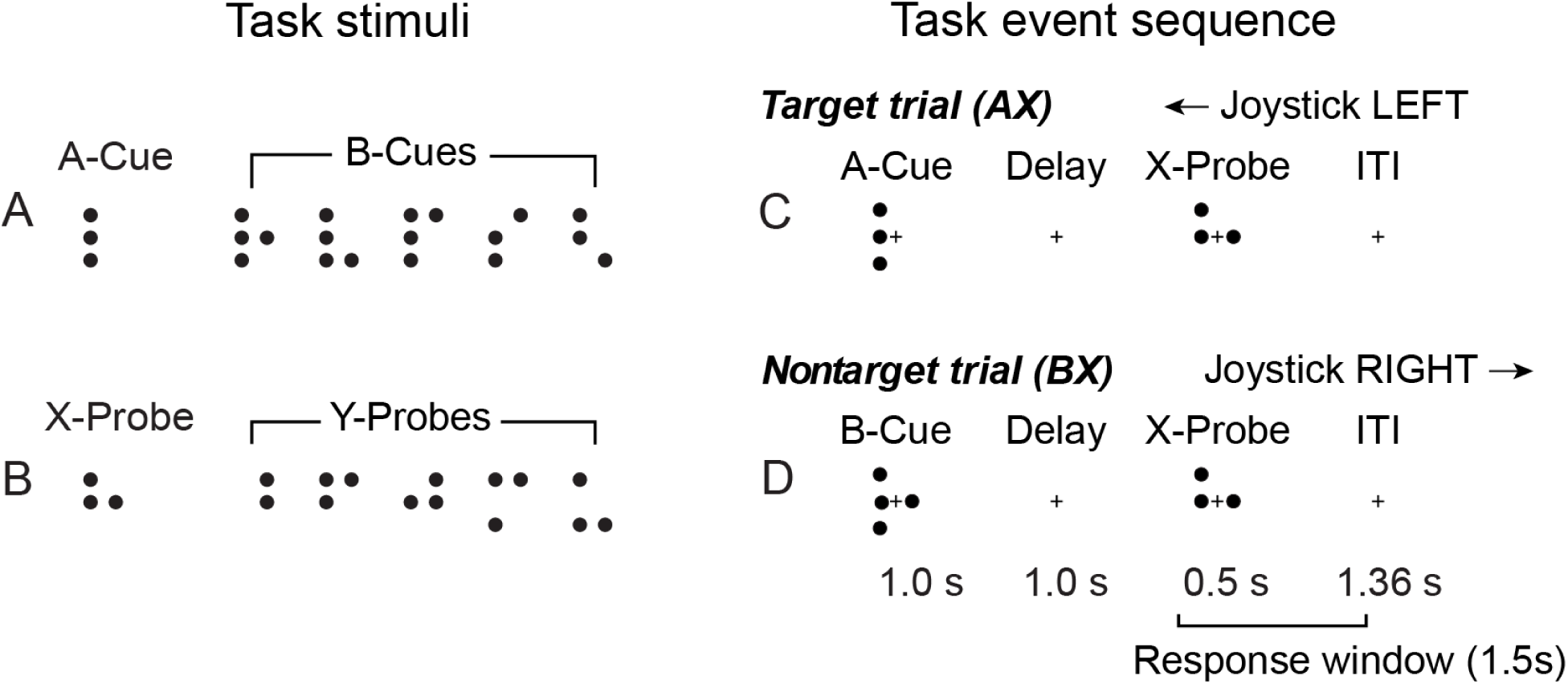
DPX stimuli and task sequence. A. Cue stimuli. Cues were either A (one dot pattern) or B (any of 5 alternative dot patterns). B. Probe stimuli. Probes were either X (one dot pattern) or Y (any of 5 alternative dot patterns). C, D. Trial events. Each trial a cue stimulus (1 s) and probe stimulus (0.5 s) were presented in sequence, separated by a delay period (1 s). Monkeys moved a joystick left or right depending on the cue-probe sequence displayed. If the A-cue was followed by the X-probe (the target sequence), monkeys were required to move the joystick left. All other cue-probe sequences (AY, BX, BY) were nontarget and required a rightward response. Thus, the same stimulus (X-probe) could require a leftward (C) or rightward (D) response, depending on the cue preceding it, requiring cognitive control. Monkeys could respond any time within 1.5 s of probe onset. C. Trial events on AX cue-probe trials. D. Trial events on BX cue-probe trials.

A typical way to model the monkey’s DPX task behavior would be to train a reinforcement learning agent to maximize the reward it receives from the environment. In this setting, the stimuli-reward relationship is one-way, i.e. mapping stimuli to responses, and the agent does not need prediction error beyond reward to learn the task. However, we propose an alternative formulation of the task: to minimize the state prediction error at each moment, incorporating predictions about sensory inputs and motor outputs (Figure 2) in addition to reward (Friston, 2010). This transforms a reward-maximization task into a sequence prediction task, and the animal’s preference for higher reward is reflected in the biased distribution of state sequences dominated by rewarded ones, i.e., synaptic weights are updated only following rewarded behavioral sequences.

**Figure 2:**
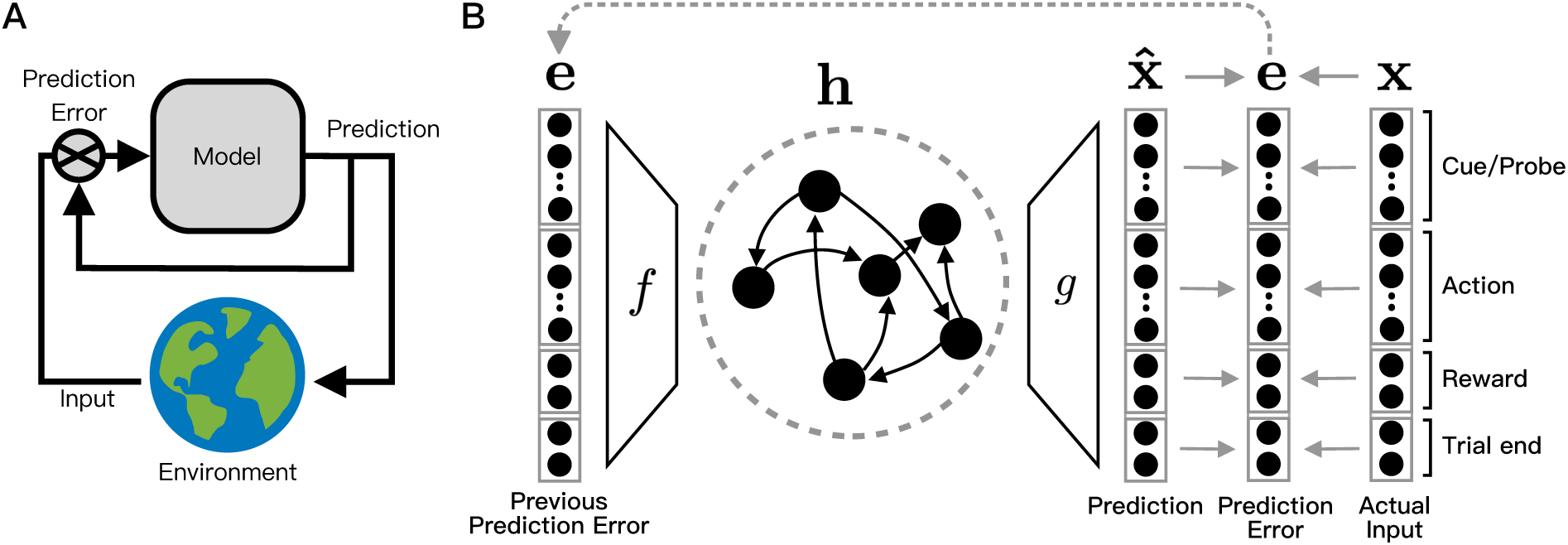
**A**: The general architecture of the model. The model predicts the environment state and utilizes its own prediction to inhibit the inputs from the environment and thus to lower the prediction error. **B**:The specific dynamics of the SPEL model. At each moment, the RNN produces a prediction 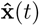 through a decoder *g*(*·*) about all incoming inputs, including cue presence/identity, probe presence/identity, response direction/occurrence, reward occurrence, and trial end. The prediction then is compared with the actual input **x**(*t*) to get an error signal in the vector form, **e**(*t*). **e**(*t*), the prediction signal, then is sent into the network via an encoder *f* (*·*). In our implementation, both *f* (*·*) and *g*(*·*) are one-layer linear projections.

With this formulation, we train a leaky recurrent neural network (RNN) to perform the DPX task Blackman et al. (2016) at a high level of proficiency (more than 90% of trials correct). To constrain the modeling approach we took inspiration from neurophysiological observations in monkey prefrontal cortex. Namely, switch neurons while not bearing a fixed relation to sensory, motor or cognitive variables, appeared to encode rare or unexpected events, namely breach in expectation. Based on this, we adapted predictive coding theory in the visual cortex (Rao and Ballard, 1999) and the principle of free energy reduction (Friston, 2010) to the training of prefrontal networks. Further, we made an architectural modification to send the residual prediction error as the input signal into the network for this RNN, which proved critical in the ability of the network to emulate neurophysiological data.

This approach is analogous to a classical closed-loop control system (Figure 2 A), in that a prediction error is fed back into the RNN as input. This contrasts with other approaches which send the relevant state of the task (not the prediction error) into the RNN at each time step as done by many similar studies (Barak, 2017; Ehrlich et al., 2021).To our knowledge, this is the first time this framework has been used to model prefrontal cortex.

Thus, the dynamics of this RNN can be written as

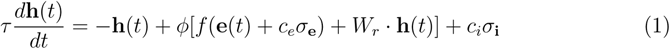

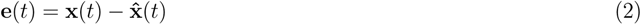

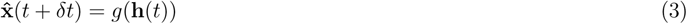

In this network (Figure 2 B), 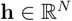 denotes the neurons in the hidden layer, where *N* is the number of neurons. The evolution of activity in hidden layer **h** is driven by its own activity through the reciprocal connectivity matrix *W_r_* and the external state prediction error **e**(*t*), both modulated by a nonlinear activation function *ϕ*. We also inject two types of independent Gaussian noises, 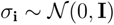 and 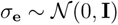, controlled by scaling coefficients *c_i_* and *c_e_*, into the equation to simulate noise from external inputs and intrinsic perturbations.

The **x** represents the state of the entire task, including at each time step in the simulation cue presence/identity, probe presence/identity, response direction/occurrence, reward occurrence, and trial end. The output of the RNN, i.e., the prediction of the next state, 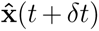, is generated by a decoder *g*, in the form of a fully connected single-layer neural network. The residual prediction error, **e**(*t*), is then fed back to the RNN through another single-layer neural network *f*. We train the RNN to generate behavioral sequences in an autoregressive way, i.e., to predict the next state 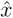 at each time step. During a training epoch, we adjust synaptic weights selectively on rewarded trials to minimize sequence prediction error (Fig. 3), after which we fix synaptic weights, and collect performance data during an evaluation epoch.

**Figure 3:**
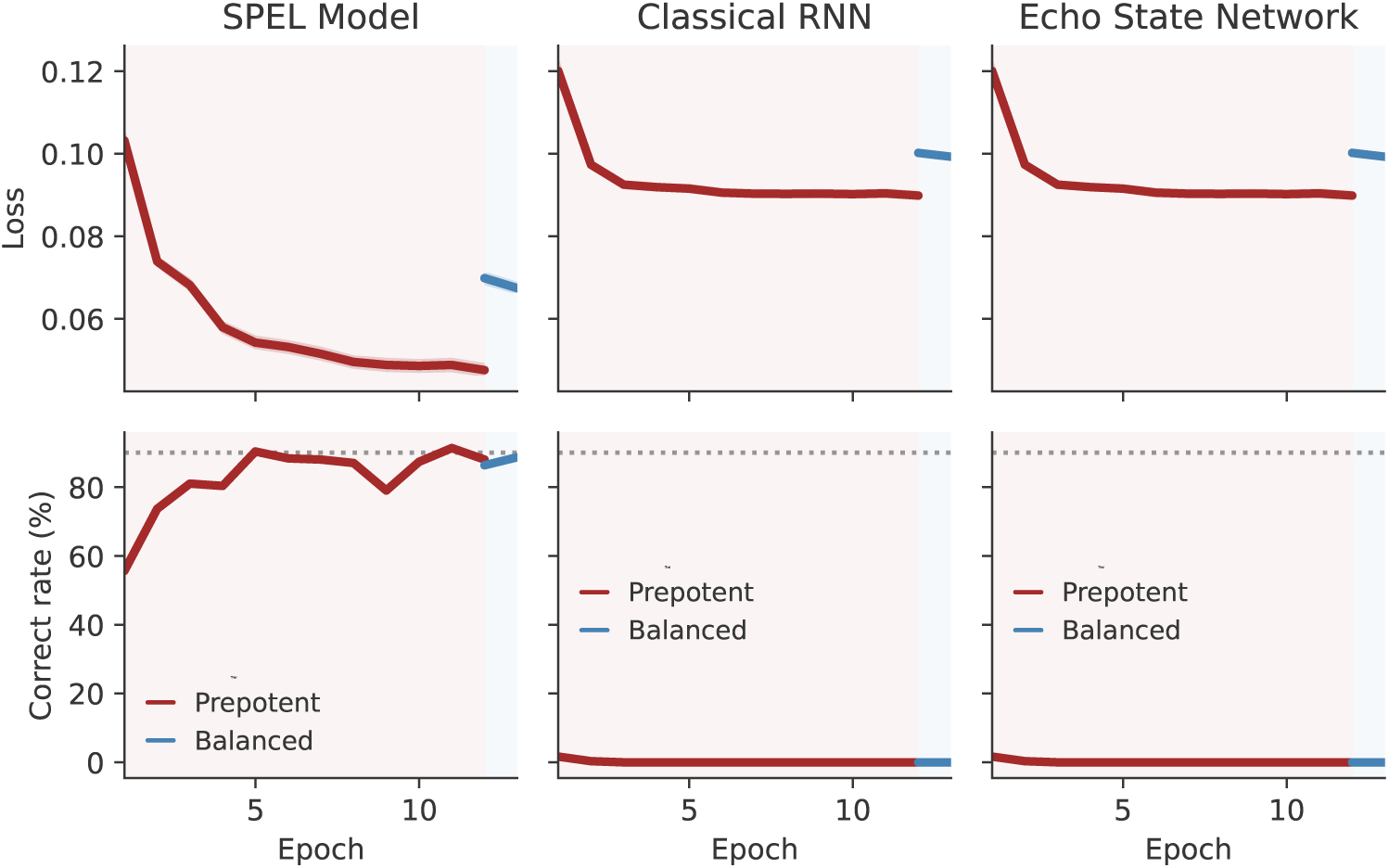
The training dynamics of the SPEL model compared with the classical RNN and the echo state network. Left: SPEL model; Middle: classical RNN; Right: echo state network (ESN). Top: Loss curve (sum squared error); Bottom: Percent correct performance. In all panels, red curves represent model performance on prepotent trial sets (in which 69% of trials presented the AX cue-probe sequence) while blue curves denote performance on balanced trial sets (in which all cue-probe combinations were presented with equal probability).

### Data analysis

#### Behavior

We consider two alternative models to the SPEL model to evaluate its performance against related models. We first test if an RNN taking raw inputs **x** rather than **e** (Figure 2 A) from the environment can learn the task. This implementation is equivalent to a classical RNN. Second, we compared the SPEL model with an echo state network (ESN) (Jaeger, 2007), to test whether error backpropagation through time is necessary. The task design, encoding paradigm, architecture, dynamics and all other hyperparameter settings of this classical RNN and the ESN are identical to the SPEL model, except that the input is **x** rather than **e** in the classical RNN and that learning in the hidden layer is deactivated in the echo state network.

The experimental results (Figure 3) demonstrate that under these conditions, the SPEL model performs as well as the classical RNN in terms of percent of trials performed correctly while maintaining stable performance when the dataset is switched from the prepotent trial set (in which 69% of trials presented the AX cue-probe sequence) to the balanced trial set (in which AX, BX, AY and BY cue-probe combinations occurred with equal frequency). However, the ESN is incapable of learning the task; the best it can achieve is learning to fixate. This suggests that the task is not so trivial that it can be solved solely by the intrinsic and fixed hidden dynamics of an RNN.

If monkey prefrontal cortex and the SPEL model acquire sequence expectations based on event frequency, response time ought to be prolonged when this expectation is broken. We found that both monkeys (Fig. 4, top) and the SPEL model (Fig. 4, bottom) exhibited longer response times (p<0.0001) to the probe on trials that the A-cue was followed by the Y-probe (Fig. 4, left). On the large majority of training trials, the A-cue was followed by the X-probe (the predominant combination across the training set). Thus presentation of the Y-probe after the A-cue was a breach of expectation. This imposed a cost in processing time, presumably needed to override the prepotent target response, and produce the nontarget response instead as required on AY trials.

**Figure 4:**
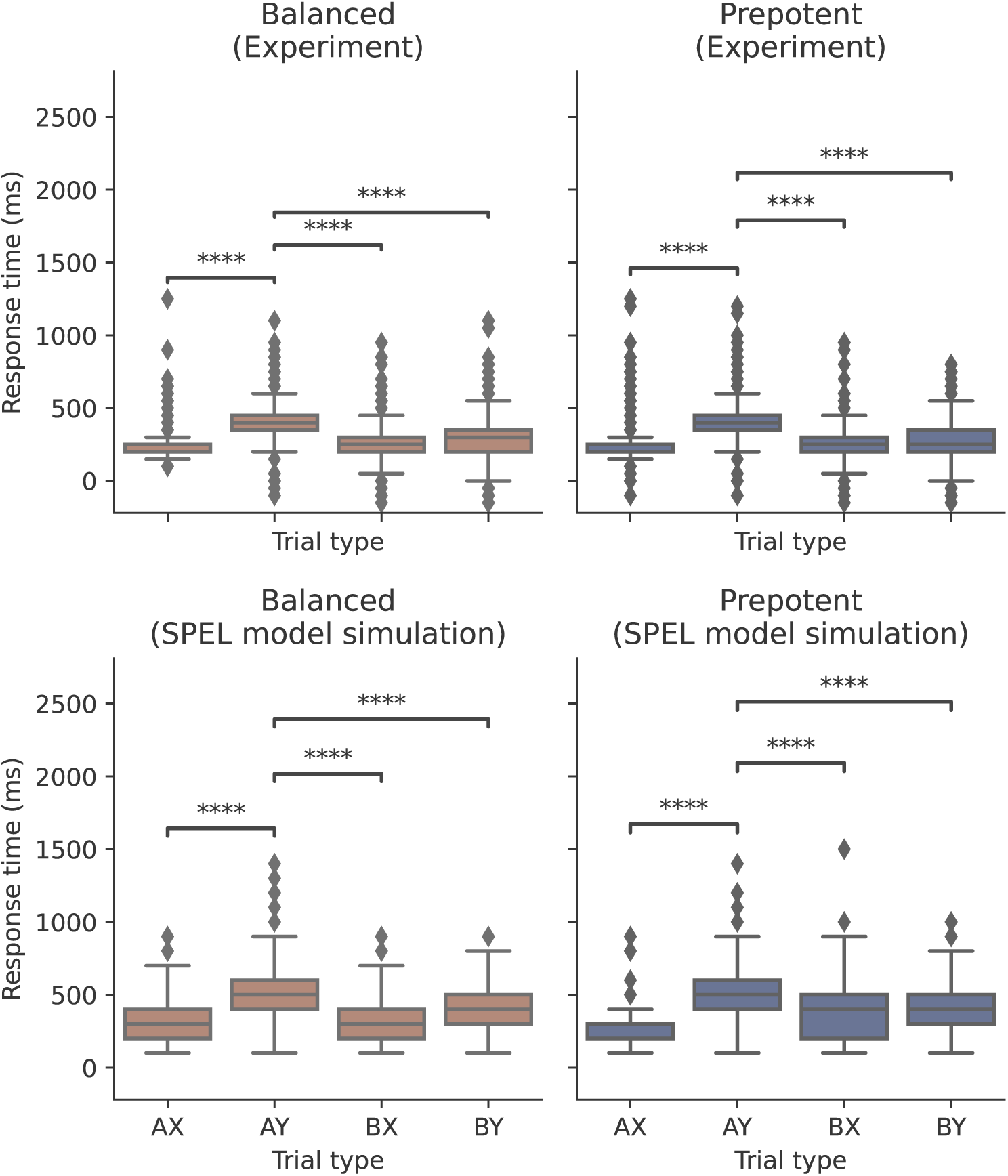
Box-whisker plots illustrate response time distributions in monkey (top) and the SPEL model (bottom) performance data during prepotent (left) and balanced (right) DPX trial sets. For both the monkey and model data, Welch’s t-tests are performed. In all conditions, the response time of AY is significantly longer than that of the other conditions (****: p <= 0.0001).

#### Neural data analysis

We used similar statistical methods to analyze the activity of 819 neurons previously recorded in monkey prefrontal cortex (Blackman et al., 2016) and 256 hidden units of the RNN trained to perform the DPX task. We first applied ANCOVA to identify units in prefrontal cortex and the RNN in which activity varied significantly as a function of the cue identity (A vs B), probe identity (X vs Y), and response (target vs nontarget). We found comparable proportions of cue, probe, and response selective neurons in monkey prefrontal cortex (Fig. 5, left) and cue, probe, and response selective units the SPEL model (Fig. 5, right). Individual neurons and units exhibited combined responses to multiple factors both in prefrontal cortex and the SPEL model. Proportionally more neurons in the SPEL model than monkey prefrontal cortex exhibited mixed selectivity for more than one DPX task variable, possibly due to the efficiency requirement of task representations in the smaller pool of units available in the SPEL model relative to prefrontal cortex.

**Figure 5:**
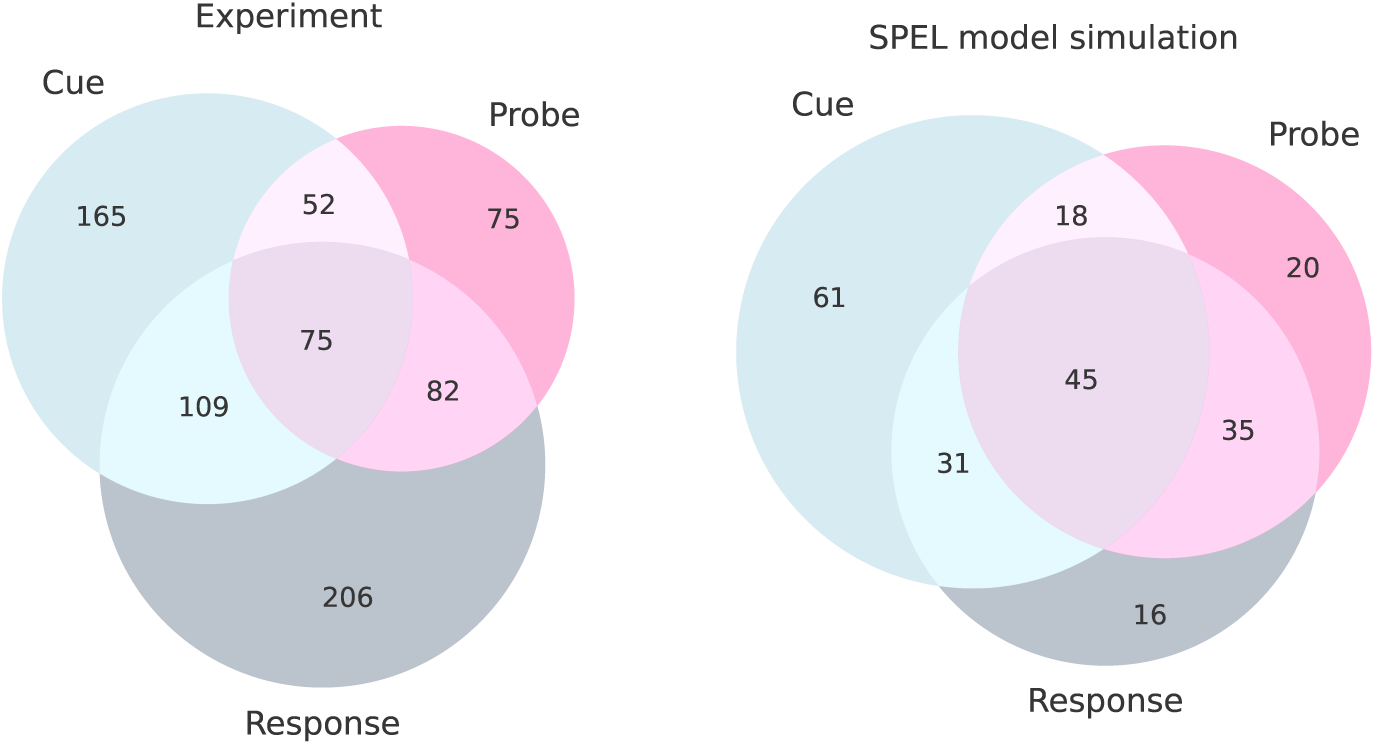
Venn diagrams illustrate the numbers of neurons in monkey prefrontal cortex (left) and the number of hidden units in the SPEL model (right), exhibiting activity that varied significantly by ANCOVA (p < 0.05) as a function of the cue (A vs B), probe (X vs Y), and response (target vs nontarget). Chi-square test shows the ratio of neurons that are selective to all variables (cue, probe, response) to those that are only selective to one variable is significantly (*p <* 0.00001) higher in the SPEL model.

We next sought to determine whether dynamical features of the SPEL model may mirror and potentially explain neural dynamics in prefrontal cortex during task performance. We addressed that question by analyzing the temporal dynamics of the mean value of the sequence prediction error vector **e**(*t*) that was used to adjust synaptic weights in the SPEL model, and that also provided input to the hidden units of the SPEL model in our architecture (Fig. 2B). We found several striking parallels between the temporal evolution of the sequence prediction error signal in the SPEL model and neural population activity in monkey prefrontal cortex. The sequence prediction error signal in the SPEL model (Fig. 6, middle) and neural activity in prefrontal cortex (Fig. 6, top) exhibited the ’switch’ pattern we previously described (Blackman et al., 2016, 2023; DeNicola et al., 2020), by which activity switched its apparent visual stimulus preference from B-cues during the cue period to A-cues during the probe period. Both the sequence prediction error signal in the SPEL model and population activity in prefrontal cortex exhibited an additional increment in activity level during the probe period when A-cues were followed by Y-probes in comparison to when A-cues were followed by X-probes (Fig. 6, top and middle, compare red and orange activity traces). Finally, these modulations in neural activity level, in both the SPEL model and in monkey prefrontal cortex, were more pronounced on prepotent trial sets (Fig. 6, top and middle, right) in comparison to balanced trial sets (Fig. 6, top and middle, left). This included augmented activity levels on B-cue trials during the cue period (Fig. 6, top and middle, black arrows) as well as augmented activity levels on AY trials during the probe period (Fig. 6, top and middle, black arrows). This is of interest in that task events associated with elevated activity levels (B-cues and AY cue-probe sequences) were less frequent, and therefore invoked larger sequence prediction errors when they occurred, in prepotent relative to balanced trial sets. Thus several idiosyncractic features of neural activity in monkey pre-frontal cortex that were difficult to account for invoking traditional correlates of prefrontal neural activity, such as selectivity for task stimuli, responses, or storage of items in working memory (Chafee and Goldman-Rakic, 1998), were readily accounted for by the SPEL model as correlates of sequence prediction error.

**Figure 6:**
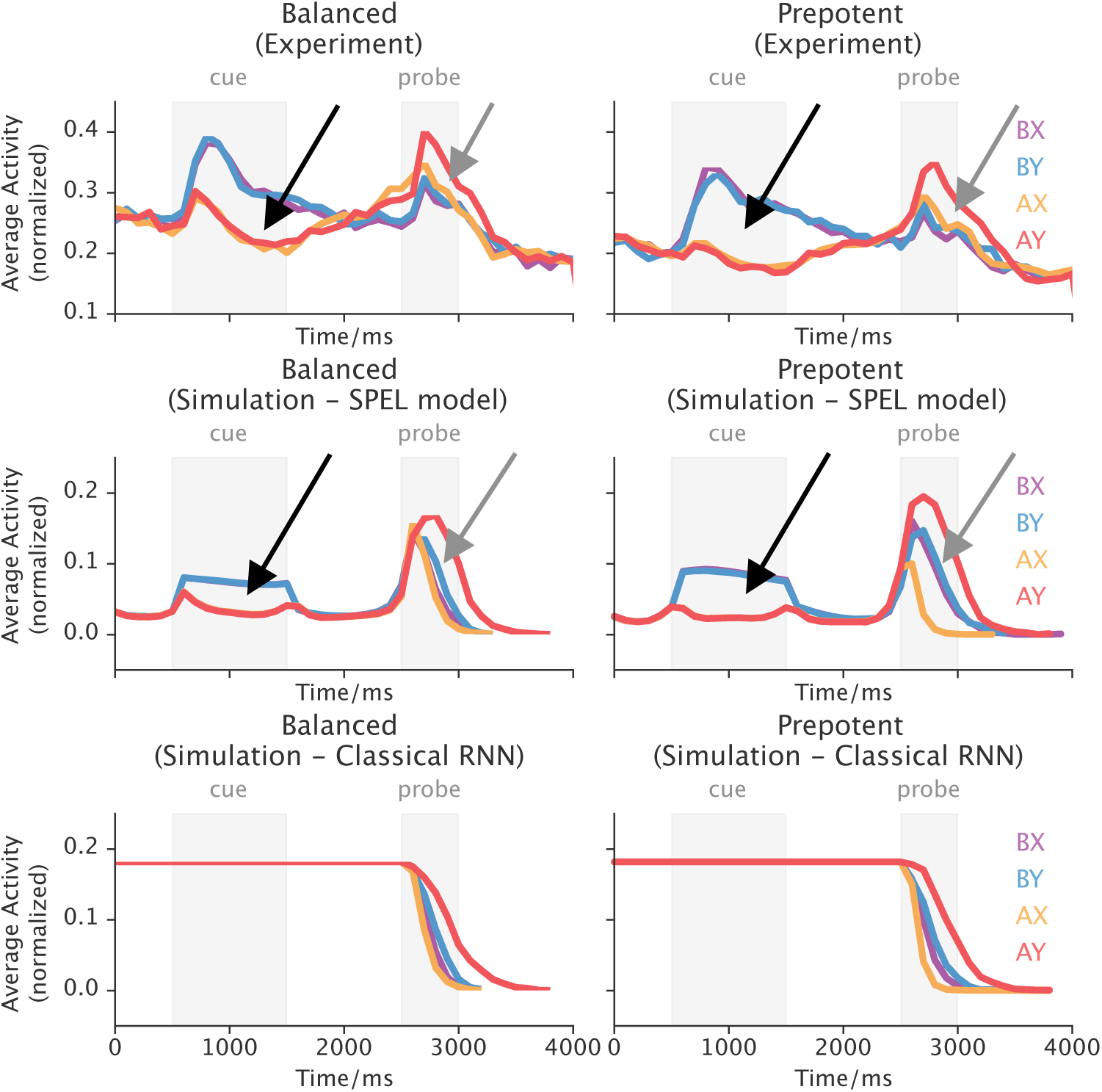
Activity dynamics in monkey prefrontal cortex (top row), the SPEL model (middle row), and a classical RNN (bottom row), during DPX task performance. Top: Neural activity in monkey prefrontal cortex exhibits the switch pattern previously described (Blackman et al., 2016, 2023; DeNicola et al., 2020). Lines plot average normalized population firing rate over time. Line color indicates cue-probe sequence (AX: orange, AY: red, BX: purple, BY: blue). Also, note that in the middle and bottom rows, some lines overlap. Activity in these neurons during the cue period is maximal when B-cues are presented (blue and purple time courses), and is maximal during the probe period when Y-probes follow A-cues (red time course). Activity on balanced trial sets is shown on the left, and on prepotent trials sets on the right (note augmentation of activity modulation in the latter case, at arrows). Middle: The average value of the input vectors, i.e., sequence prediction error vector in the previous time step, in the SPEL model (conventions as in the top panels) as a function of time within the trial. Note comparability of activity dynamics, including augmentation of modulation on Prepotent trial sets (arrows), in the SPEL model in comparison to monkey prefrontal cortex. Bottom: Input activity dynamics in a classical RNN. For most of the time the **average** input activity is flat due to the aforementioned encoding design. Note discrepancy of neural dynamics in the classical RNN and in monkey prefrontal cortex.

Equally of interest, these results were not generic to recurrent neural networks trained to perform the DPX task. The classical RNN trained as rapidly and reached similar asymptotic performance levels as the SPEL model when performing the DPX task (Figure 3). Yet, the input neurons in the classical RNN failed to replicate the neural activity dynamics observed in prefrontal cortex entirely (bottom row in Fig.6). This suggests that alternative architectures and learning rules can achieve a desired level of performance on a given task, but only some of these learning rules and architectures may be implemented in prefrontal cortex.

We had previously reported that the neural representation of the probe stimulus in pre-frontal cortex was contingent on the identity of the preceding cue, according to the logical constraints of the DPX task. Specifically, prefrontal neurons effectively encoded the identity of the probe only on trials when the preceding cue was A (Blackman et al., 2016). This is the only case in which the identity of the probe mattered, from the perspective of having an impact on behavior, since on B-cue trials, the identity of the probe (X or Y) was irrelevant to the response required, which was nontarget in all cases. The selective encoding of probe identity when it bore logical relation to task performance provided, in addition to switch activity, a unique neural signature in prefrontal cortex of computations required to perform the DPX task. We found that the neural representation of the probe by hidden units in the SPEL model was similarly dependent on the identity of the probe. We decoded probe identity (X vs Y) from patterns of activity in monkey prefrontal cortex (Fig. 7, top), and hidden units within the SPEL model (Fig. 7, bottom) within a sliding window passed through the trial. Decoding accuracy in both prefrontal cortex and the SPEL model climbed from chance levels to 60-70% decoding accuracy shortly after probe stimulus onset in the case that the preceding cue had been A (Fig. 7, top and middle, left, pink trace), but not in the case that the preceding cue had been B (Fig. 7, top and middle, right, gray trace). Note the peak decoding accuracy in the monkey neural data is lower than that of Blackman et al. (2023) as the temporal resolution is decreased in this study as we aligned the electrophysiological data with the model data, while the general trend is the consistent in both studies. Thus the SPEL model, like the prefrontal cortex, only encoded sensory information when it was useful for predicting upcoming events. This result is particularly noteworthy because previous cognitive models trained on a very similar task have considered whether some learned gating is necessary to control what is and is not stored in working memory (Hazy et al., 2006), and indeed this motivated the LSTM model (Hochreiter and Schmidhuber, 1997) as well as the attention control found in transformers (Vaswani et al., 2017). Our results show that for the purposes of learning what information to encode into prefrontal networks, such gating can be learned via existing recurrent neural network mechanisms, without need of additional gating mechanisms.

**Figure 7:**
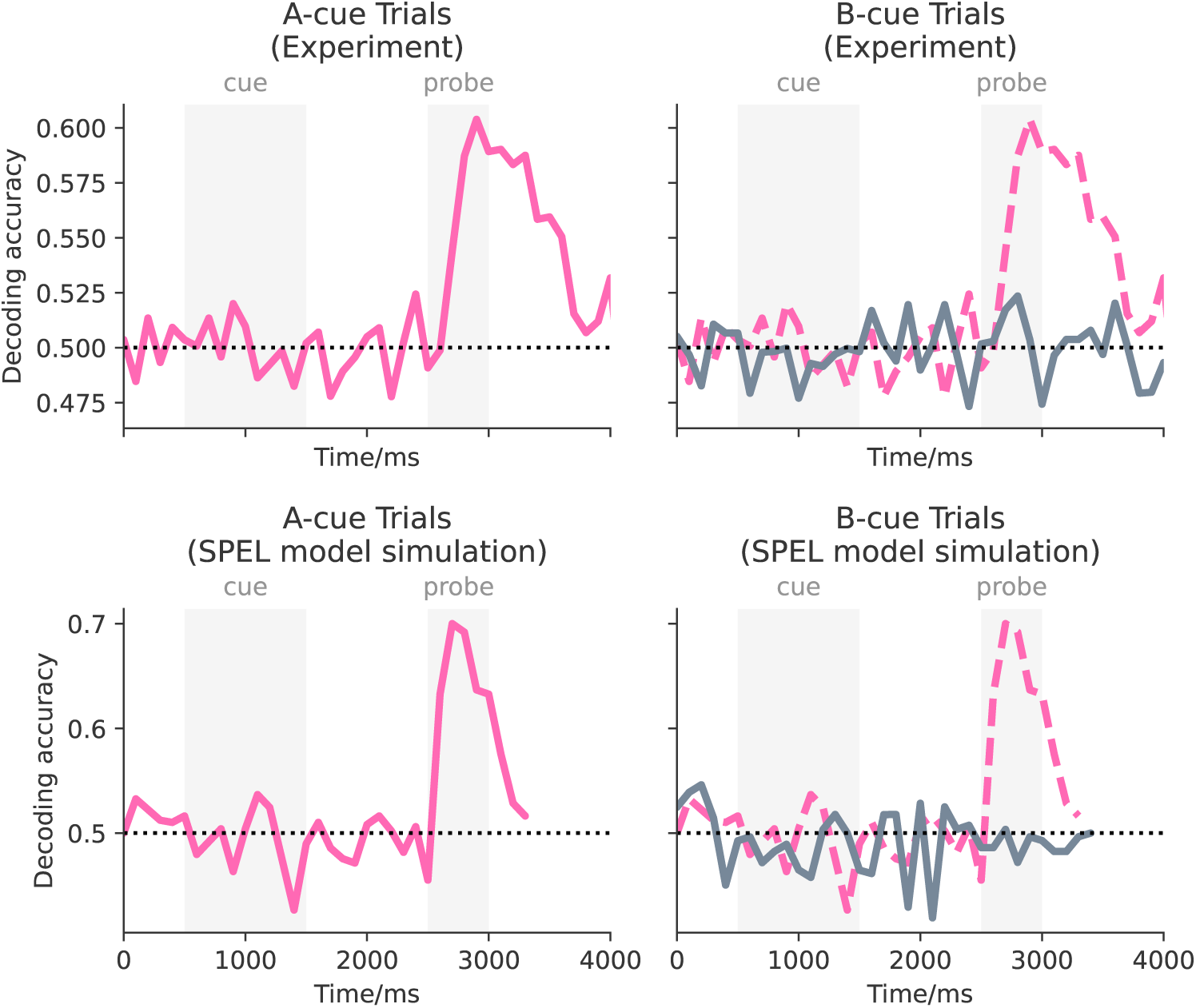
Cue-dependent representation of probe information in prefrontal cortex and the SPEL model. Time courses plot the accuracy with which the probe (x vs Y) was decoded by a linear classifier applied to activity patterns in monkey prefrontal cortex (Top) and in hidden units of the SPEL model (bottom) within a sliding window passed through the trial. Probe decoding accuracy is plotted separately on trials that the preceding cue was A (left, pink time course) and B (right, gray time course - dashed pink time course on A-cue trials superimposed for comparison).

## 3 Discussion

We have shown above that a recurrent neural network model that simply predicts the next state on the basis of prediction errors can provide a detailed account of neurophysiological effects in prefrontal cortex that are otherwise of unclear origin (Blackman et al., 2016, 2023; DeNicola et al., 2020). The SPEL model follows previous applications of deep networks to vision (Kriegeskorte, 2015), demonstrating that deep networks can also model the lateral prefrontal cortex. Previous models of prefrontal cortex have included various rate coded (Alexander and Brown, 2015; Hazy et al., 2006) and spiking (Compte et al., 2000; Calvin and Redish, 2021) approaches, which share common features of relatively complex architectures and some need for hand tuning of parameters. Spiking models have generally not simulated the ways in which representations are formed, although they have simulated biophysical details such as the dynamics of NMDA channels. Previous lateral PFC models have also not captured error activity in lateral PFC, although some previous models have simulated error-related activity in medial PFC (Alexander and Brown, 2018, 2011). Some previous models have also explored explicit gating signals to control working memory updating (Hazy et al., 2006), and this may not be necessary given that representations are stored only as needed, as shown in Figure 7, as an indirect consequence of the learning rule we applied (minimizing sequence prediction error). The SPEL model is essentially a standard recurrent network with few free hyperparameters and structural degrees of freedom, which suggests that its ability to capture neurophysiological data and representations is not merely curve fitting.

Empirically, prior neural recording studies have demonstrated prefrontal neurons encode reward prediction error (RPE) (Oemisch et al., 2019; Voloh et al., 2020), as implemented by classical temporal difference reinforcement learning models. Prior modeling studies of prefrontal cortex have used RPE to train networks that capture certain aspects of neural activity (Alexander and Womelsdorf, 2021). Reward prediction error signals are, like sequence prediction error signals, a difference between predictions and outcomes, but in the case of RPE the outcomes are driven by actions and their consequences (reward), rather than by the continuously evolving temporal structure of the environment. To our knowledge no prior modeling study has attempted to generate artificial neural signals approximating those recorded in prefrontal cortex using a recurrent architecture trained in a sequence prediction error learning framework. The close parallels between biology and theory readily achieved using this approach has strong implications for the nature of the computations performed by prefrontal cortex, and the neural mechanisms by which these computations emerge through learning. Our results suggest that prefrontal cortex automatically learns the temporal structure of the environment by encoding sequence prediction error signals into firing rates directly, in essence becoming a self-teaching network.

The SPEL model somewhat uniquely uses its own prediction error signals both as teaching signals and as inputs to the model. With backpropagation through time as a training signal, this provides a convenient reduction of the delta rule training signal – instead of requiring both the input and error terms, the model can simply use the error terms from two consecutive time steps as a training signal, which may simplify the computations. Furthermore, we have recently shown that backpropagation through time can be closely approximated by a biologically plausible learning rule that is purely local in space and time (Cheng and Brown, 2023), and that performs full gradient learning. This suggests that the SPEL model can be adapted in a straightforward manner as a biologically plausible model.

The SPEL model currently has several limitations. First, the hidden units show a strong pattern of temporal oscillation, with a period that depends highly on the time constant *τ*, more so than we see in monkey single units which generally show more sustained activity. This suggests that the model found a different solution for hidden unit representation compared with the macaque brain. Nonetheless, hidden units in the SPEL model exhibited conditional encoding of task information, as did prefrontal cortex (Fig. 7). Second, we have approximated the receptive fields of input neurons with one-hot codes. We anticipate that the difference will matter little in practice, but a more realistic model would use Gaussian rather than one-hot receptive fields for inputs. Third, the model error neuron activities are computed simply as the difference between actual states (*X*) and predicted states (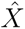), rather than feeding this difference into a governing equation as in equation (1). Again, we anticipate this will make little difference in practice.

In sum, our results indicate that a relatively simple RNN trained to perform a cognitive control task by minimizing sequence prediction error generates neural and behavioral dynamics that bear striking resemblance to behavioral dynamics in monkeys and neural dynamics observed in monkey prefrontal cortex. Moreover, this was not a trivial result in that these dynamical features of the experimental data were only captured by some and not other RNN implementations, according to the nature of the learning rule implemented. This has equally important implications for both biological and computational investigations of prefrontal cortex. From a biological perspective, it is of deep interest what utility is achieved by prefrontal neurons encoding a training signal into firing rates explicitly, literally representing the error in the firing rates of neurons. It may be that this makes it possible for activity patterns in prefrontal cortex to govern activity-dependent forms of synaptic plasticity (Feldman, 2012; Dan and Poo, 2004), making it possible for prefrontal cortex to serve as its own teacher. From a computational perspective, our results suggest that RNNs trained to perform cognitive tasks by minimizing sequence prediction error offer a comparatively simple, generalized computational framework in which a single architecture using a single learning rule may be trained to perform a variety cognitive tasks in a way that mimics the computational strategy of prefrontal cortical networks.

## 4 Methods

### Nonhuman primate behavioral and neurophysiological methods

Neural data were recorded in prefrontal cortex of monkeys performing a cognitive control task and were described in prior reports (Blackman et al., 2023, 2016). We briefly describe behavioral and neurophysiological methods here.

#### Subjects

We recorded neural activity in the dorsolateral prefrontal cortex of two monkeys performing the dot-pattern expectancy (DPX) task (Blackman et al., 2023, 2016; Jones et al., 2010)(Figure 1A-D). The DPX task is a variant of the widely-used AX-CPT in which letters are replaced with dot-pattern (Jones et al., 2010). All animal care and experimental procedures complied with National Institutes of Health guidelines and were approved by the Animal Care and Use Committee at the University of Minnesota and Minneapolis Veterans Administration Medical Center.

#### Behavioral Paradigm

The DPX Task: Monkeys maintained gaze fixated on a central target while a cue followed by a probe stimulus was presented each trial. One dot pattern was designated the A-cue, whereas five alternative dot patterns were collectively designated B-cues (Fig. 1C). Similarly, one probe dot pattern was designated the X-probe, whereas five probe dot patterns were collectively designated Y-probes (Fig. 1D). After the cue-probe sequence, monkeys moved a joystick to the left or right. The rewarded response direction depended on the cue-probe sequence presented. If the cue was A and probe was X (AX sequence, target trial), the correct response direction was to move the joystick to the left. All other cue-probe combinations (nontarget trials) required a rightward joystick response. We administered the DPX task in both balanced and prepotent trial sets. In prepotent sets, the four possible cue-probe combinations were equiprobable (AX, AY, BX, BY). In prepotent trial sets, the majority of trials presented the target AX cue-probe sequence (69%) whereas the remaining minority of trials (31%) presented nontarget cue-probe sequences (AY 12.5%, BX 12.5%, BY 6%).

#### Neural recording

To record the spiking activity of individual prefrontal neurons, we advanced 16 independently movable electrodes into the prefrontal cortex. Spike waveforms of individual neurons were isolated online (Alpha Omega Engineering, Alpharetta, GA, BAK Electronics, Umatilla, FL). Time stamps of action potential detection and task events were saved to disk for offline analysis.

### Model methods

#### RNN implementation

The RNN is implemented using PyTorch (Paszke et al., 2019), following the discretization (Equation 1):

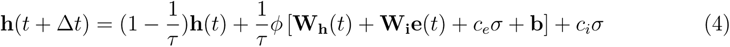

where *ϕ* is set to tanh, Δ*t* is set to 100 ms, *τ* is set to 5, *c_i_* is set to 0.25, *c_e_* is set to 0.5, and *σ* is set to multivariate standard Gaussian noise with mean zero and diagonal variance one. **W_i_**, **W_i_** and **b** are corresponding input and recurrent weight matrices with biases. Based on Equation 4, the prediction of the task state 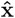 is generated by 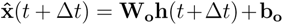, where **W_o_** is the output weight matrix and **b_o_** is its bias. Specifically, **x** in Equation 4 is a 22-d vector representing all cognitive variables related to the DPX task, shown in Figure 2. The first 12 dimensions represent the 12 possible visual inputs to the RNN, corresponding to 12 distinct visual patterns for cues and probes. The next 6 dimensions model the animal’s response, formed by combinations of two types of actions: fixations (FIXATION, NO-FIXATION) and joystick movements (LEFT, CENTER, RIGHT). The following 2 components form a one-hot vector representing the presence of rewards. The final 2 components form another one-hot vector indicating whether the current trial has terminated.

#### Training details

In our experiments, the dimensionality *N* of 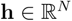, denoted as the number of neurons in the hidden layer, is set to 256. The SPEL model is optimized with an Adam optimizer (Kingma and Ba, 2014) set at a learning rate of 0.0005 with betas of (0.9, 0.999). The loss function is the mean squared error (MSE) between the output of the RNN, **x**(*t*) = **W***_o_*(*t*), and the input, 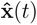. The *ℓ*_2_ norm of the weights is penalized during training with a decay factor of 0.0005. During training, for each epoch, we randomly generate mature behavior data, encode them into sequences of 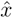 and use them as datasets to optimize the RNN in an autoregressive way, like the pretraining stage in other sequence predictions such as language models (Radford et al., 2018). The RNN converges after approximately 12 epochs on the prepotent trial dataset during training. It is subsequently retrained on the balanced trial dataset and achieves a similar correctness rate in just 2 epochs, given that the task rule remains unchanged. Post each epoch, the RNN’s performance is evaluated on a simulated DPX environment as a traditional RL agent, wherein the model’s action is generated by the 12th-18th component of **x** within the RNN itself. The training is halted when the RNN’s performance on the simulated environment reaches a threshold of 90% accuracy or when the maximal training epoch is reached.

For the comparison with the classical RNN and echo state network (Figure 3), all the parameters for the training and model architecture are the same, except for the necessary changes in gradient descent for the echo state network and the input form changes from **e** to **x** in the classical RNN case.

### Data analysis

The data analysis is done similarly to Blackman et al. (2016), with slight modifications to accommodate both the model data and monkey data. First, we preprocess both data into firing rate sequences with a bin size of 100ms. Next, neurons in the model and monkey data were nonexclusively categorized into three types (Figure 5) using the ANCOVA test, with a p-value threshold of 0.0001. To obtain the decoding accuracy for cue types (Figure 7), a linear regression model was utilized. The score was obtained using cross-validation with this model. The software used for all statistical analyses above were scikit-learn (Pedregosa et al., 2011) and Pingouin (Vallat, 2018).

### Code availability

The code needed to reproduce all the results can be found here: https://github.com/chenghuzi/SPE

## Acknowledgements

This neural data reported in this study was supported by the National Institutes of Health R01MH077779, R01MH107491, and P50MH119569; the Department of Veterans Affairs; the American Brain Sciences Chair; the Wilfred Wetzel Graduate Fellowship; and National Institute of General Medical Sciences T32 GM008244 and T32 HD007151. This work was performed while R.K.B. was employed at the University of Minnesota. The opinions expressed in this article are the author’s own and do not reflect the views of the U.S. Food and Drug Administration, the Department of Health and Human Services, or the U.S. Government. This material is the result of work supported with resources and the use of facilities at the Minneapolis VA Health Care System. The contents do not represent the views of the U.S. Department of Veterans Affairs, the National Institutes of Health, the Department of Health and Human Services, or the U.S. Government. We thank Sofia Sakellaridi and Adele DeNicola for excellent assistance with neural recording, C. Dean Evans for technical assistance during surgeries and neural recordings as well as exemplary animal care, and Dale Boeff for assistance with computer programming as well as design and construction of neurophysiological recording equipment.

## Notes

### Competing Interest Statement

The authors have declared no competing interest.

## References

Alexander, W. H. and J. W. Brown (2011, September). Medial prefrontal cortex as an action-outcome predictor. Nat. Neurosci. 14 (10), 1338–1344.

Alexander, W. H. and J. W. Brown (2015). Hierarchical Error Representation: A Computational Model of Anterior Cingulate and Dorsolateral Prefrontal Cortex. Neural Computation, 1–57.

Alexander, W. H. and J. W. Brown (2018, December). Frontal cortex function as derived from hierarchical predictive coding. Scientific Reports 8 (1), 3843–3843.

Alexander, W. H. and T. Womelsdorf (2021, February). Interactions of medial and lateral prefrontal cortex in hierarchical predictive coding. Front. Comput. Neurosci. 15, 605271.

Averbeck, B. B., J.-W. Sohn, and D. Lee (2006, February). Activity in prefrontal cortex during dynamic selection of action sequences. Nat. Neurosci. 9 (2), 276–282.

Barak, O. (2017). Recurrent neural networks as versatile tools of neuroscience research. Current opinion in neurobiology 46, 1–6.

Barch, D. M., C. S. Carter, A. W. MacDonald, III, T. S. Braver, and J. D. Cohen (2003, February). Context-processing deficits in schizophrenia: Diagnostic specificity, 4-week course, and relationships to clinical symptoms. J. Abnorm. Psychol. 112 (1), 132–143.

Barch, D. M. and A. Ceaser (2012). Cognition in schizophrenia: core psychological and neural mechanisms. Trends Cogn. Sci. 16 (1), 27–34. doi: 10.1016/j.tics.2011.11.015. Epub 2011 Dec 12.

Bellet, M. E., M. Gay, J. Bellet, B. Jarraya, S. Dehaene, T. van Kerkoerle, and T. I. Panagiotaropoulos (2021, October). Spontaneously emerging internal models of visual sequences combine abstract and event-specific information in the prefrontal cortex.

Blackman, R. K., D. A. Crowe, A. L. DeNicola, S. Sakellaridi, A. W. MacDonald, 3rd, and M. V. Chafee (2016, April). Monkey prefrontal neurons reflect logical operations for cognitive control in a variant of the AX continuous performance task (AX-CPT). J. Neurosci. 36 (14), 4067–4079.

Blackman, R. K., D. A. Crowe, A. L. DeNicola, S. Sakellaridi, J. A. Westerberg, A. M. Huynh, A. W. MacDonald, S. R. Sponheim, and M. V. Chafee (2023, April). Shared neural activity but distinct neural dynamics for cognitive control in monkey prefrontal and parietal cortex. J. Neurosci. 43 (15), 2767–2781.

Brown, J. W., D. Bullock, and S. Grossberg (2004, May). How laminar frontal cortex and basal ganglia circuits interact to control planned and reactive saccades. Neural Netw. 17 (4), 471–510.

Calvin, O. L. and A. D. Redish (2021, May). Global disruption in excitation-inhibition balance can cause localized network dysfunction and schizophrenia-like context-integration deficits. PLoS Comput. Biol. 17 (5), e1008985.

Chafee, M. V. and P. S. Goldman-Rakic (1998). Matching patterns of activity in primate prefrontal area 8a and parietal area 7ip neurons during a spatial working memory task. J. Neurophysiol. 79 (6), 2919–2940.

Chen, L. L. and S. P. Wise (1995, March). Neuronal activity in the supplementary eye field during acquisition of conditional oculomotor associations. J. Neurophysiol. 73 (3), 1101–1121.

Cheng, H. and J. W. Brown (2023, February). Replay as a basis for backpropagation through time in the brain.

Compte, A., N. Brunel, P. S. Goldman-Rakic, and X. J. Wang (2000, September). Synaptic mechanisms and network dynamics underlying spatial working memory in a cortical network model. Cereb. Cortex 10 (9), 910–923.

Crowe, D. A., S. J. Goodwin, R. K. Blackman, S. Sakellaridi, S. R. Sponheim, A. W. MacDonald, 3rd, and M. V. Chafee (2013, October). Prefrontal neurons transmit signals to parietal neurons that reflect executive control of cognition. Nat. Neurosci. 16 (10), 1484–1491.

Dan, Y. and M.-M. Poo (2004, September). Spike timing-dependent plasticity of neural circuits. Neuron 44 (1), 23–30.

DeNicola, A. L., M.-Y. Park, D. A. Crowe, A. W. MacDonald, 3rd, and M. V. Chafee (2020, February). Differential roles of mediodorsal nucleus of the thalamus and prefrontal cortex in Decision-Making and state representation in a cognitive control task measuring deficits in schizophrenia. J. Neurosci. 40 (8), 1650–1667.

Ehrlich, D. B., J. T. Stone, D. Brandfonbrener, A. Atanasov, and J. D. Murray (2021). Psychrnn: an accessible and flexible python package for training recurrent neural network models on cognitive tasks. eneuro 8 (1).

Feldman, D. E. (2012, August). The spike-timing dependence of plasticity. Neuron 75 (4), 556–571.

Freedman, D. J., M. Riesenhuber, T. Poggio, and E. K. Miller (2002, August). Visual categorization and the primate prefrontal cortex: neurophysiology and behavior. J. Neurophysiol. 88 (2), 929–941.

Friston, K. (2010, February). The free-energy principle: a unified brain theory? Nat. Rev. Neurosci. 11 (2), 127–138.

Friston, K., J. Mattout, and J. Kilner (2011, February). Action understanding and active inference. Biol. Cybern. 104 (1-2), 137–160.

Gläscher, J., N. Daw, P. Dayan, and J. P. O’Doherty (2010, May). States versus rewards: dissociable neural prediction error signals underlying model-based and model-free reinforcement learning. Neuron 66 (4), 585–595.

Goldman-Rakic, P. S. (1987, December). Circuitry of primate prefrontal cortex and regulation of behavior by representational memory. https://onlinelibrary.wiley.com › doi › cphy.cp010509 https://onlinelibrary.wiley.com › *doi › cphy.cp010509*, 373–417.

Goodwin, S. J., R. K. Blackman, S. Sakellaridi, and M. V. Chafee (2012, March). Executive control over cognition: stronger and earlier rule-based modulation of spatial category signals in prefrontal cortex relative to parietal cortex. J. Neurosci. 32 (10), 3499–3515.

Hazy, T. E., M. J. Frank, and R. C. O’Reilly (2006, April). Banishing the homunculus: making working memory work. Neuroscience 139 (1), 105–118.

Hochreiter, S. and J. Schmidhuber (1997, November). Long short-term memory. Neural Comput. 9 (8), 1735–1780.

Jaeger, H. (2007). Echo state network. scholarpedia 2 (9), 2330.

Jones, J. A. H., S. R. Sponheim, and A. W. MacDonald, 3rd (2010, March). The dot pattern expectancy task: reliability and replication of deficits in schizophrenia. Psychol. Assess. 22 (1), 131–141.

Kingma, D. P. and J. Ba (2014). Adam: A method for stochastic optimization. *arXiv preprint arXiv:1412.6980*.

Kriegeskorte, N. (2015, November). Deep neural networks: A new framework for modeling biological vision and brain information processing. Annu. Rev. Vis. Sci. 1 (1), 417–446.

Lesh, T. A., A. J. Westphal, T. A. Niendam, J. H. Yoon, M. J. Minzenberg, J. D. Ragland, M. Solomon, and C. S. Carter (2013, April). Proactive and reactive cognitive control and dorsolateral prefrontal cortex dysfunction in first episode schizophrenia. Neuroimage Clin 2, 590–599.

MacDonald, 3rd, A. W. (2008, July). Building a clinically relevant cognitive task: case study of the AX paradigm. Schizophr. Bull. 34 (4), 619–628.

MacDonald, 3rd, A. W., C. S. Carter, J. G. Kerns, S. Ursu, D. M. Barch, A. J. Holmes, V. A. Stenger, and J. D. Cohen (2005, March). Specificity of prefrontal dysfunction and context processing deficits to schizophrenia in never-medicated patients with first-episode psychosis. Am. J. Psychiatry 162 (3), 475–484.

McCloskey, M. and N. J. Cohen (1989). Catastrophic interference in connectionist networks: The sequential learning problem. Volume 24 of Psychology of Learning and Motivation, pp. 109–165. Academic Press.

Miller, E. K. and J. D. Cohen (2001). An integrative theory of prefrontal cortex function. Annu. Rev. Neurosci. 24, 167–202.

Nieder, A. and E. K. Miller (2003, January). Coding of cognitive magnitude: compressed scaling of numerical information in the primate prefrontal cortex. Neuron 37 (1), 149–157.

Oemisch, M., S. Westendorff, M. Azimi, S. A. Hassani, S. Ardid, P. Tiesinga, and T. Womelsdorf (2019, January). Feature-specific prediction errors and surprise across macaque fronto-striatal circuits. Nat. Commun. 10 (1), 176.

Paszke, A., S. Gross, F. Massa, A. Lerer, J. Bradbury, G. Chanan, T. Killeen, Z. Lin, N. Gimelshein, L. Antiga, et al. (2019). Pytorch: An imperative style, high-performance deep learning library. Advances in neural information processing systems 32.

Pedregosa, F., G. Varoquaux, A. Gramfort, V. Michel, B. Thirion, O. Grisel, M. Blondel, P. Prettenhofer, R. Weiss, V. Dubourg, et al. (2011). Scikit-learn: Machine learning in python. the Journal of machine Learning research 12, 2825–2830.

Radford, A., K. Narasimhan, T. Salimans, I. Sutskever, et al. (2018). Improving language understanding by generative pre-training.

Ramirez-Cardenas, A., M. Moskaleva, and A. Nieder (2016, May). Neuronal representation of numerosity zero in the primate parieto-frontal number network. Curr. Biol. 26 (10), 1285–1294.

Rao, R. P. and D. H. Ballard (1999). Predictive coding in the visual cortex: a functional interpretation of some extra-classical receptive-field effects. Nature neuroscience 2 (1), 79–87.

Rigotti, M., O. Barak, M. R. Warden, X.-J. Wang, N. D. Daw, E. K. Miller, and S. Fusi (2013, May). The importance of mixed selectivity in complex cognitive tasks. Nature.

Vallat, R. (2018). Pingouin: statistics in python. J. Open Source Softw. 3 (31), 1026.

Vaswani, A., N. Shazeer, N. Parmar, J. Uszkoreit, L. Jones, A. N. Gomez, L. u. Kaiser, and I. Polosukhin (2017). Attention is all you need. In I. Guyon, U. V. Luxburg, S. Bengio, H. Wallach, R. Fergus, S. Vishwanathan, and R. Garnett (Eds.), Advances in Neural Information Processing Systems, Volume 30. Curran Associates, Inc.

Voloh, B., M. Oemisch, and T. Womelsdorf (2020, September). Phase of firing coding of learning variables across the fronto-striatal network during feature-based learning. Nat. Commun. 11 (1), 4669.

Wallis, J. D., K. C. Anderson, and E. K. Miller (2001). Single neurons in prefrontal cortex encode abstract rules. Nature 411 (6840), 953–956.

Zarr, N. and J. W. Brown (2023, January). Foundations of human spatial problem solving. Sci. Rep. 13 (1), 1485.

